# Modeling the Tangled Bank: A Users Guide to Gillespie Eco-evolutionary Models using the Julia Programming Language

**DOI:** 10.1101/2025.10.13.682111

**Authors:** Dipanjana Dalui, John P DeLong

## Abstract

1. Eco-evolutionary processes drive patterns in species’ abundances and traits. However, modeling complex ecological systems is challenging because of the many species and traits involved and because processes unfold in stochastic and non-equilibrium conditions. Gillespie Eco-evolutionary Models (GEMs) were created to simulate the eco-evolutinary dynamics in such complex scenarios.
2. GEMs bridge the gap between ecological and evolutionary timescales, enabling tracking of trait dynamics and abundances simultaneously, and allowing real-time feedback between the two processes. Designed to capture demographic stochasticity and individual trait variation, GEMs enable tracking of multi-trait and multi-species evolution. Originally designed for a quantitative genetics approach, GEMs have now been expanded to handle both quantitative traits and discrete traits (genotypes or strains).
3. In this manuscript, we introduce an optimized and accessible framework for GEMs developed in the Julia programming language. We have simplified the simulation setup process by placing many of the information containers and design structures into functions that adapt to the model configurations chosen by the user. We use a birth-death logistic growth model as an example to illustrate the GEM setup.
4. This *user’s guide* is designed to make building GEMs for new models more streamlined and approachable. In addition to compartmentalized internal workings, the new framework also improves computational speed and efficiency. This lets users focus on model development rather than algorithmic complexities.

## 1 Introduction

Ecological systems are extremely complex, with numerous interacting species and myriad dynamical processes influencing their structure and function. Ecological systems emerge from short- and long-term eco-evolutionary processes that generate patterns of abundances and traits across species (Wickman et al. 2024, DeLong 2020). Moreover, ecological systems are subject to stochasticity and may often be in non-equilibrium states (Shoemaker et al. 2020, Hastings et al. 2018, Kummel and Vasseur 2025), creating complexity that can be difficult to understand. This complexity was the backdrop of Darwin’s musing about evolution in a “tangled bank” (Darwin 1859).

Models of ecological systems are often grounded in the dynamics of a single species or of simple food web modules for tractability (Polis and Winemiller 2013). However, several approaches have emerged that capture eco-evolutionary dynamics and provide insights into complex patterns of trait and abundance change over time. These approaches include quantitative genetics (Lande 1979, Abrams et al. 1993, McPeek 2017), adaptive dynamics (Geritz et al. 1998), integral projection models (Ellner and Rees 2006, Metcalf et al. 2016), and individual agent-based models (Mollet et al. 2016, Enberg et al. 2009, Ayllón et al. 2016). Despite these advances, it is still challenging to capture eco-evolutionary dynamics in complex ecological scenarios with multiple interacting species wherein many traits may be under selection in stochastic and non-equilibrium conditions (i.e., tangled banks). A more recent approach - Gillespie eco-evolutionary model (GEM) simulations - was created to enable eco-evolutionary modeling in such complex tangled-bank scenarios.

Like other eco-evolutionary modeling approaches, GEMs also unify the ecological and evolutionary timescales to track the dynamics of traits and abundances simultaneously, allowing for real-time feedback between the two processes. This feedback is essential for understanding complex ecological systems. Historically, evolution has been thought to operate on a longer timescale than ecological dynamics (Schoener et al. 2002), but a growing body of theory and data shows that evolutionary change can occur rapidly enough to impact - and be impacted by - ecological processes as they happen (DeLong et al. 2016, Hairston Jr et al. 2005, Palkovacs and Hendry 2010). GEMs achieve this eco-evolutionary feedback (or feedforward) for an - in principle - unlimited number of interacting species in non-equilibrium and stochastic conditions (DeLong and Belmaker 2019, DeLong and Luhring 2018, DeLong and Coblentz 2022).

In addition to being able to handle complex scenarios, there are at least three additional benefits of modeling eco-evolutionary dynamics with GEMs. First, GEMs are built on the Gillespie stochastic simulation algorithm (SSA) (Gillespie 1977, 2007), which uses a stochastic birth-and-death process to generate change in population abundances over time. The Gillespie SSA already has broad use in ecology (Boettiger 2018; Kummel and Vasseur 2025; Yaari et al. 2012). Building GEMs on the Gillespie SSA means that demographic stochasticity is always present in a GEM simulation. The effects of stochasticity can then play out in the simulations, being minimal in large populations but generating more drift-like dynamics in smaller populations. Furthermore, because individuals vary in traits (see detailed explanations below), stochasticity can create differences among individuals that supersede the fitness value of the trait, generating individual demographic stochasticity (Van Daalen and Caswell 2017).

Second, GEMs do not use a fixed level of trait variation. Heritable trait variation is the raw material for evolution. Not only does it determine a population’s potential to evolve, it also plays an important role in shaping ecological interactions with potentially important ecological consequences (Bolnick et al. 2011; Schreiber et al. 2011; Vasseur et al. 2011). Despite this, many existing models either ignore trait variation altogether (Yoshida et al. 2003) or treat it as static (Schreiber et al. 2011; Vasseur et al. 2011), failing to capture its dynamic nature and its interplay with ecological interactions. In GEMs, trait variation changes with each gain or loss of an individual in the population, fluidly responding to and influencing subsequent dynamics.

Third, fitness functions - equations relating traits to fitness - are not needed for GEMs. This advantage becomes increasingly important as the system being modeled becomes more complex. Most eco-evolutionary model approaches require an *a priori* mathematical link between traits and fitness (for example, see Schreiber et al. 2011 or Vasseur et al. 2011), which requires assumptions about how parameters control demography, often using the Breeder’s equation (Abrams et al. 1993; Heywood 2005). Although fitness functions lend themselves to analytical understanding of models, they may make strong assumptions about the underlying biology. In addition, fitness functions can, in some cases, impose fixed fitness landscapes that do not reflect changing ecological conditions, even if the links between fitness and traits do not stay constant over time (Siepielski et al. 2009).

Despite the availability of some GEM code in online repositories, an introductory resource on how GEMs work and how to begin to use them is lacking, potentially making the uptake of this tool challenging. A deeper understanding of the mathematical nature of the Gillespie SSA and GEMs has been covered elsewhere (Coblentz and DeLong 2023, Kummel and Vasseur 2025). In this manuscript, we introduce and describe an upgraded, accessible, and significantly optimized framework for GEMs. The current version - developed in Julia - provides computational speed and efficiency and compartmentalizes much of the internal workings so that users can focus on model development. In this document, we aim to demonstrate how this new computational tool can simplify entry into eco-evolutionary modeling with GEMs, enabling researchers to model real-time feedback between these dynamical processes while accounting for heritable trait variations and bypassing the need for a fitness function. We hope that this will enable the scientific community to undertake more complex eco-evolutionary simulations that more closely approach the workings of Darwin’s tangled bank. With this manuscript, we aim to facilitate a deeper understanding of eco-evolutionary dynamics by providing a flexible and powerful modeling platform that addresses key limitations of existing approaches. To keep the algorithm accessible for adaptation, and encourage versatility in its research application, we are publishing the GEMs as a set of Julia scripts, with the capacity to be loaded as a local package. We do not wish to make accessing and modifying the core algorithm code harder for other researchers.

In the following sections, we will introduce the Julia-based GEM framework in detail and provide a comprehensive guide to its implementation. In section 2, we will detail the algorithm’s framework and then walk the readers through the steps of setting up a GEM in section 3. This will include step-by-step help in defining the model, creating the birth-and-death terms, and configuring the parameter space. Section 4 will cover running the GEM simulation, accessing the simulation output, and visualization. And finally, we cover some advance considerations like running GEMs in parallel and using GEMs as a local package in section 5.

## 2 What are GEMs

The Gillespie algorithm is a multi-step state-dependent Poisson process that simulates demo-graphic stochasticity at an individual level and generates trajectories of finite populations in continuous time. For any given population of *N* individuals at time *t*, a birth occurring with probability *b*(*N*(*t*)) will increase the population by 1 (*N* → *N* + 1), and a death occurring with probability *d*(*N*(*t*)) will decrease the population by 1 (*N* → *N* − 1). These independent birth- and-death events are Poisson processes. The change in the probability of being in state *N* is given by the sum of the ways to enter the state, minus the ways to leave the state, which is the master equation for an expression of probability balance. The “propensity” of each possible event is defined as the rate of the event divided by the sum of all event rates. The probability of selecting an event is proportional to its propensity. Time is then advanced by a value *τ* drawn from a logistic distribution with mean defined by the sum of all rates *S*

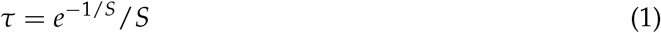

*τ* and the total abundance across all populations,*N*_*tot*_, are inversely related, meaning that more events occur per unit time as population densities increase as a result of increased encounters with between individuals.

Gillespie Eco-evolutionary Models (GEMs), first introduced by DeLong and Gibert 2016, work by allowing changes to be driven by direct but stochastic fitness consequences of traits. That is, the traits possessed by individuals (traits may have equal parameters) determine the propensities of birth and death in the context of the model, and these propensities then guide the demographic consequences for individuals, and thus the whole population. As the simulation of the model proceeds, a sequence of births and deaths occurs. In the event of a birth, the GEM calculates the offspring trait using the parent’s trait value, the population trait distribution, and the heritability of this trait. This offspring trait is now added to the population’s trait distribution. In the event of a death, that individual (with the corresponding trait values) is deleted from the population’s trait distribution. Over time, birth and death events tend to amplify trait values that offer a fitness advantage (high likelihood of birth, low likelihood of death). As a result, the final trait distribution is centered around more ecologically fit values. And as individuals are gained and lost through time, trait distributions update, changing population size and trait distributions together through time.

## 3 Setting up GEMs

### 3.1 Getting Started

All scripts for running GEMs are located in the GitHub repository at (https://github.com/dipanjanadalui/Julia-GEM). The Julia-GEM framework we introduce requires Julia Programming Language v1.10 (Bezanson et al. 2017) or higher to run. We recommend using an Integrated Development Environment (IDE) like VSCode to keep the workflow organized. The GitHub repository also hosts example models, a quick start guide, and regularly updated content. To keep the workflow simple, scripts that are unlikely to be modified by a new user are stored in the `functions` directory. When setting up a GEM analysis, new users will need to interact with only the files for model definition (`model_definition.jl`) and model configuration (`model_config.jl`), and run the analysis either interactively using the Read-Eval-Print-Loop (REPL) or through the provided `GEM_run.jl` file. Figure 1 is a visual representation of the workflow setup.

**Figure 1:**
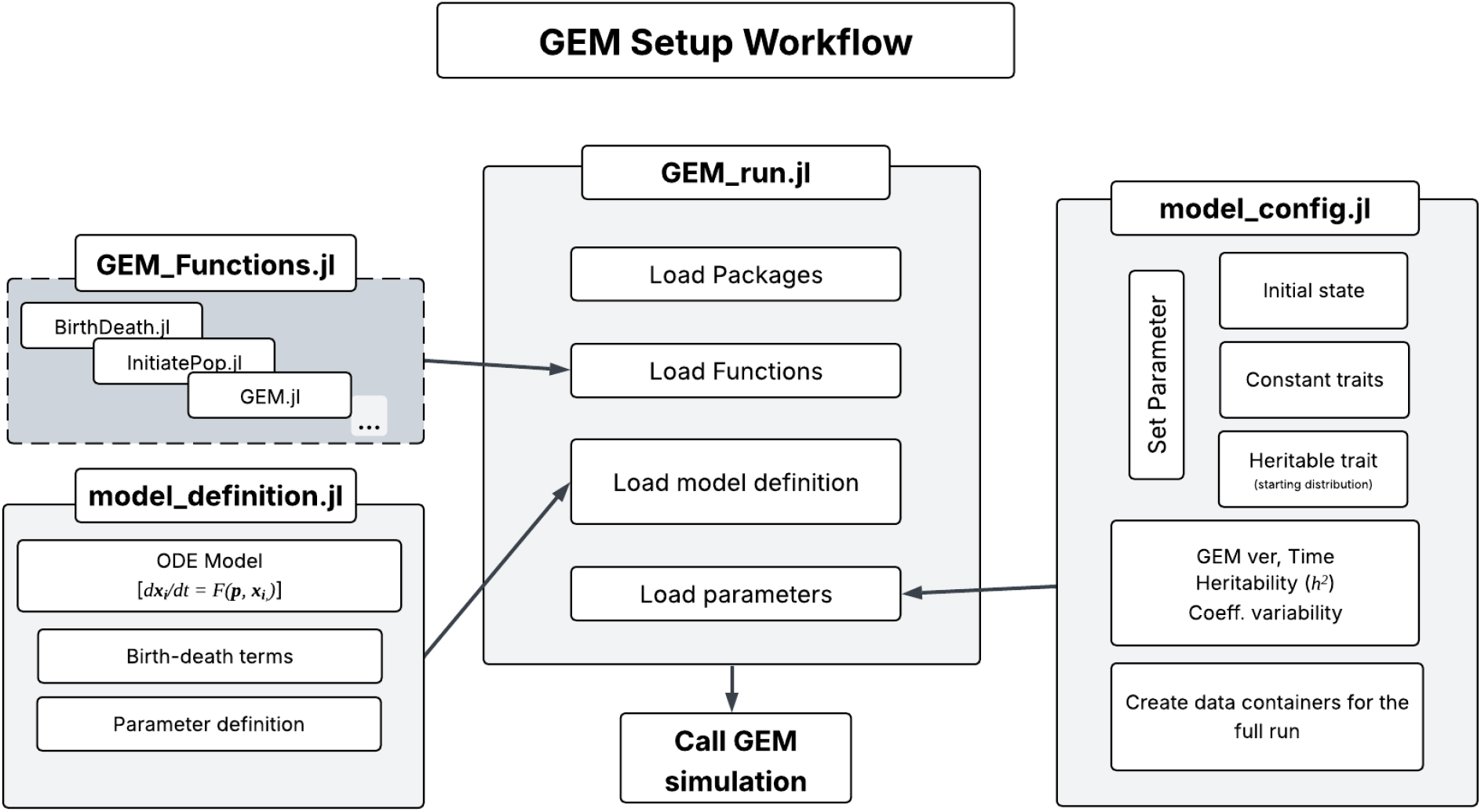
A graphical representation of the GEM setup workflow. The two scripts user will spend most of their time on while setting up new model analysis are model_definition.jl and model_config.jl. Any new functions can be either included directly in the GEM_run.jl script, or added to the list of function scripts in the GEM_functions.jl file for repeated use.

**Figure 2:**
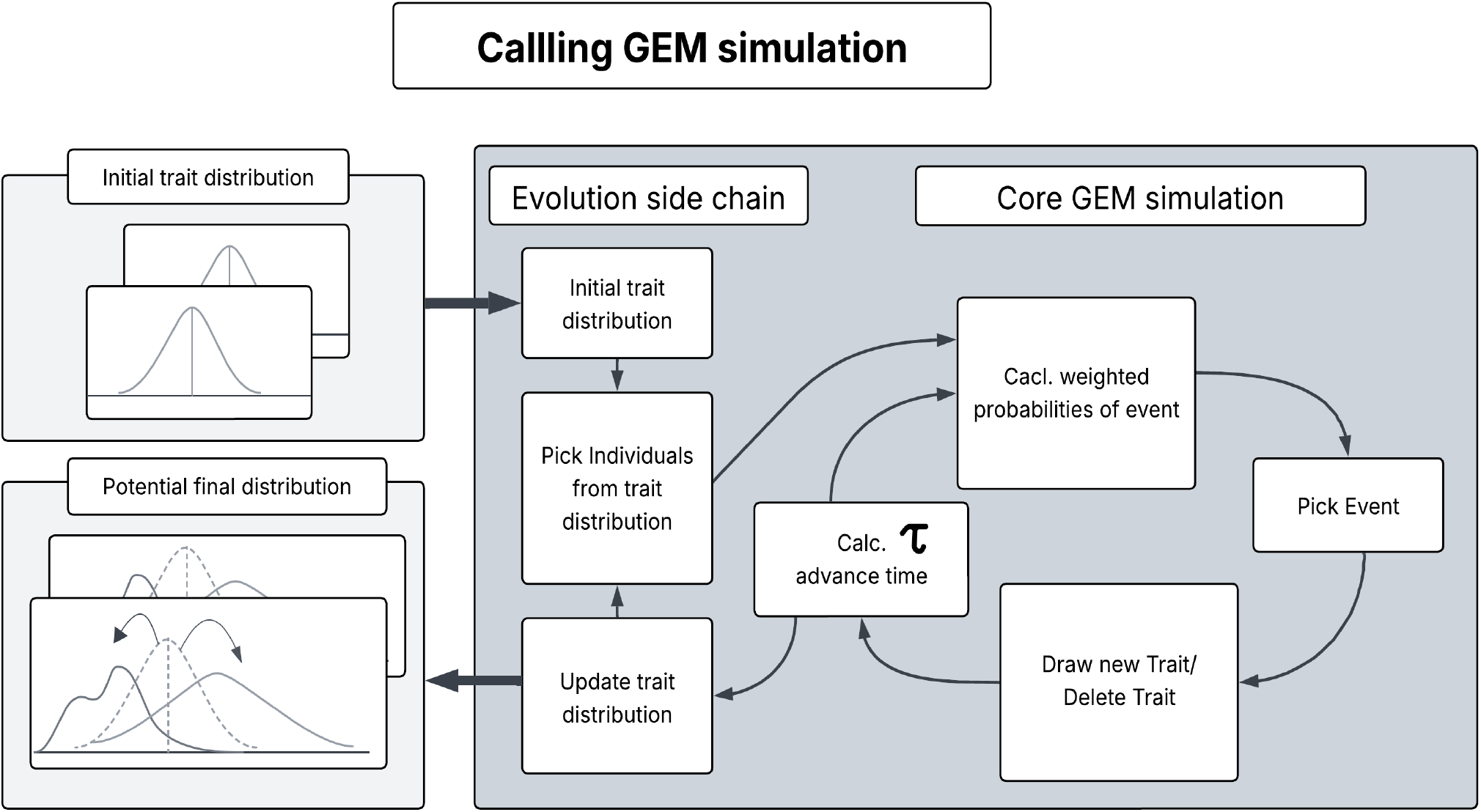
A schematic of the core GEM function with an evolutionary side chain.

The core files are

`model_definition.jl`: this file is used to define and organize the model parameters. Additionally, the custom birth-death function for the model is also defined in this script. New model structures also should be defined in this file.

`model_config.jl`: this file is used to instantiate the model’s parameters using starting values and initial state abundances.

`GEM_run.jl`: this file loads all the required packages and functions. It serves as a primary script for calling GEM functions, saving simulation data, and generating plots.

In the following sections we outline the essential components of GEMs that a new user will customize to their eco-evolutionary modeling needs. We will illustrate the concepts using the birth-death logistic model (henceforth, b-d logistic model, DeLong and Cressler 2023) as an example. A two-state (i.e., with two populations) GEM example can also be found in the GitHub repository.

### 3.2 Model Definition file

In the model definition file, there are three blocks of code to set up the ODE model itself, break out birth and death event terms, and define model parameters.

#### 3.2.1 Model

The first step involves defining the ordinary differential equation (ODE) model that describes the dynamics of the system. In general, it is expressed as

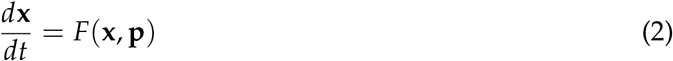

where **x**_**i**_ = {*x*_1_, *x*_2_, …, *x*_*n*_} represents the states (population, species, age-structures, etc.) and **p** = {*p*_1_, *p*_2_, …, *p*_3_} defines the model parameters. GEMs have no upper limit on the number of ODEs that you can include in your model, although increasing complexity will come at a cost in run time.

For the the case of a logistic model of the form,

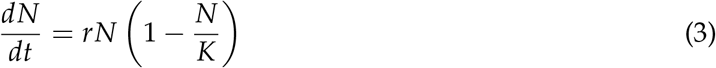

the birth-death equivalent would be

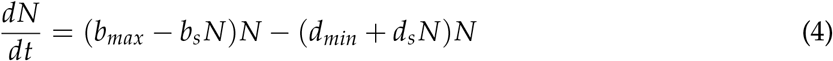

where

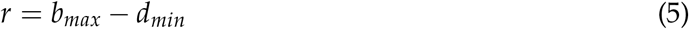

and

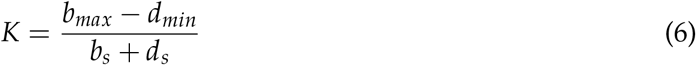

The Julia code will look something like this –

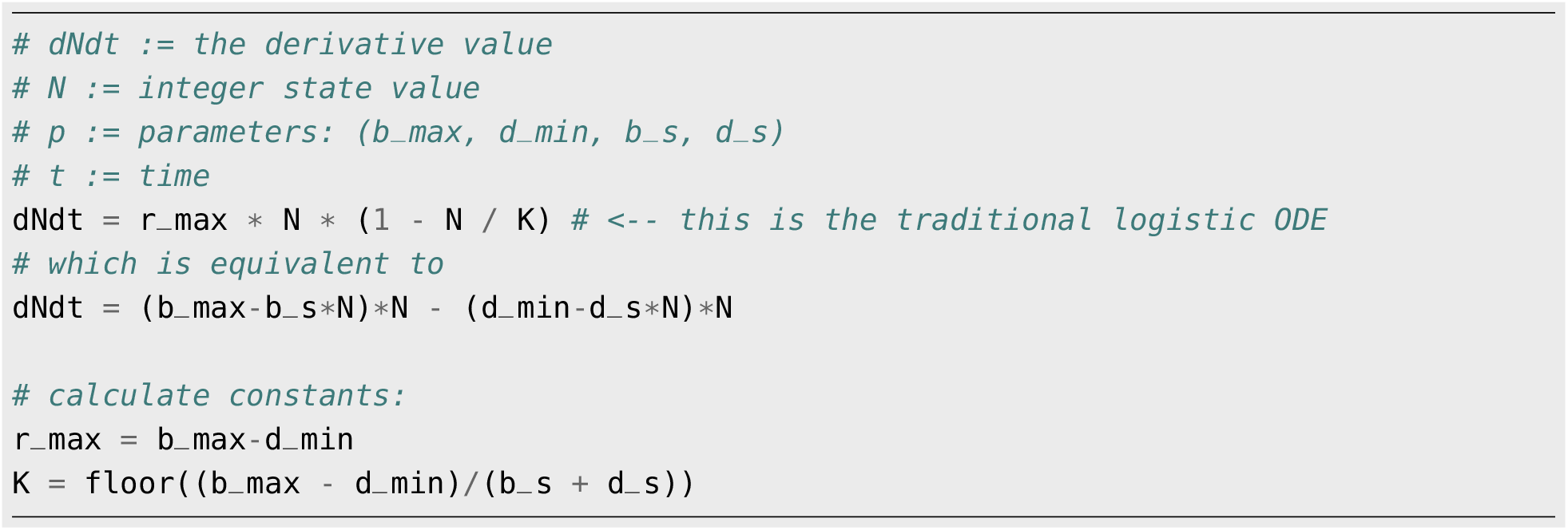

#### 3.2.2 birth-death Event Terms

In this block, the birth and death functions for each state are defined. Because the birth-death process must separate birth events from death events, intrinsic population growth rate parameters must be decomposed into separate birth and death components, which is why we use the bd-logistic rather than the standard logistic model. In addition, the effects of competition, predation, and all other interactions should also be considered in the birth-and-death functions for each state.

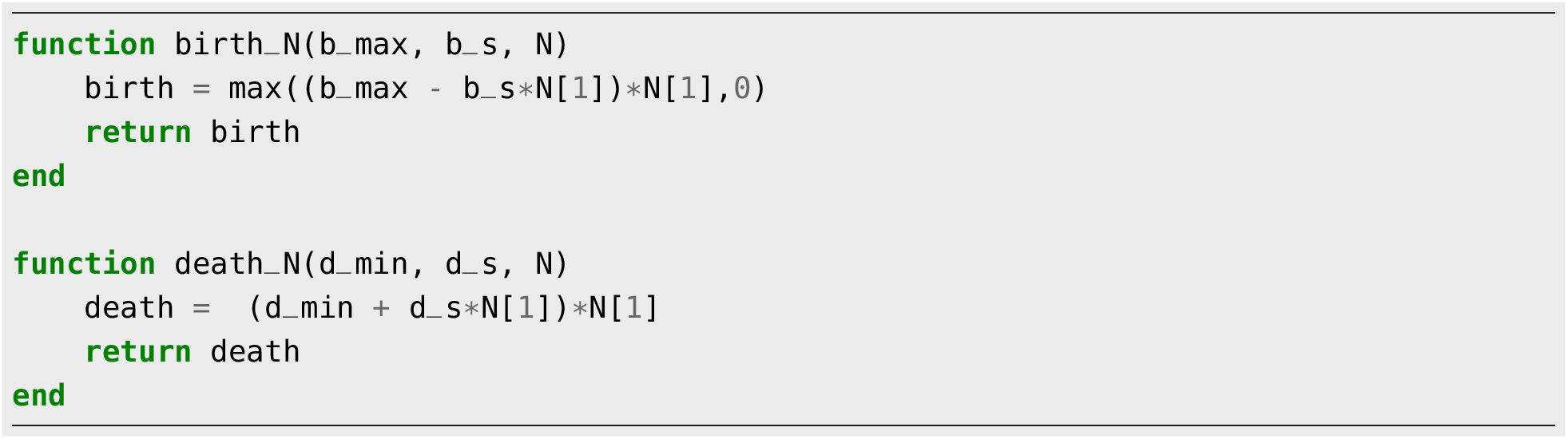

The function event_terms is defined in the same block. This function is used to generate events within the GEM algorithm that are then randomly sampled.

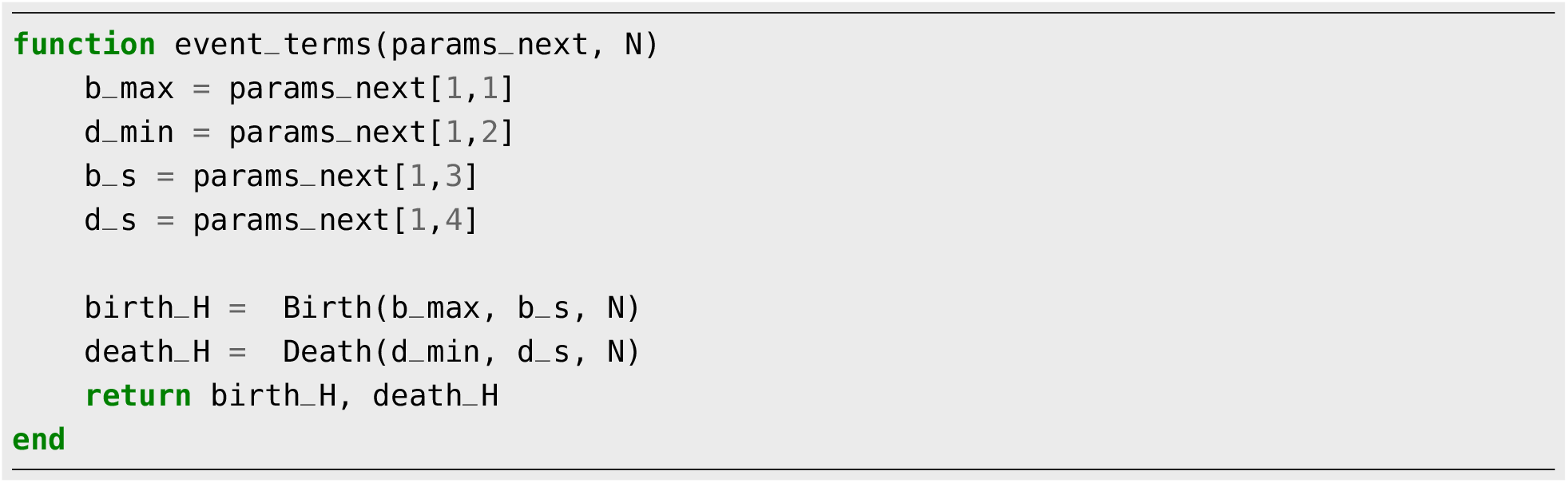

For an event to be calculated correctly as the simulation progresses, the parameter names must be correctly mapped to their corresponding values. This can be accomplished by matching the dimensions of `param_next` to the dimensions of `state_par_match` (see section 3.3.2). The row index corresponds to the state and the column index to the parameter. This means that we matching the names of the parameters for each state with a 1 in `state_par_match`. In a onestate model such as the bd-logistic, all parameters are mapped onto state N, but in a multi-state model (e.g., predator-prey), the other states (such as the predator) would get 0s rather than 1s for b_max, d_min, etc.

#### 3.2.3 Parameter Definitions

The last block of this script is used to organize the parameters for the model and the simulation into custom structures (`struct`). Similar parameters and variables are collected in `struct` that are passed to functions internally within the GEM simulation (table 1). These structures are permanent once defined. Unless a new `struct` is added, there is no need to edit this block.

**Table 1:**
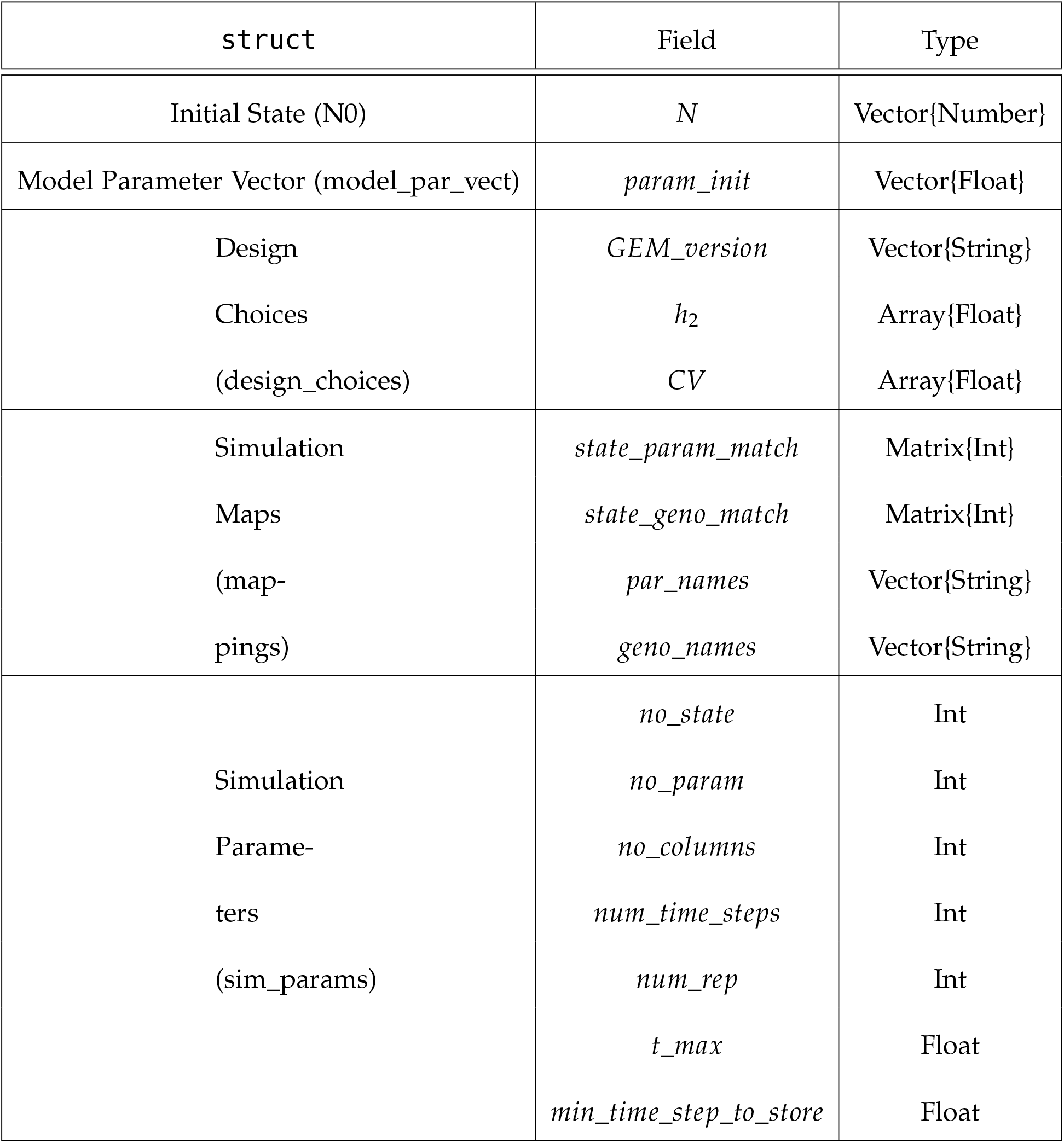
Table of struct, their fields (objects defined in the struct), and type. Names inside parenthesis in the struct column correspond to the GEM_sim function arguments.

### 3.3 Model Configuration file

Once the model is fully defined, it is time to choose initial conditions, values for the model parameters, and simulation parameters. All of these steps happen in the model configuration file.

#### 3.3.1 Initial state and parameters

This block is often where one will spend most of the time making decisions. Here, we choose an initial abundance for each state and configure the parameter space. For parameters, begin by defining a mean (*µ*) value for each parameter – this would be the traditional parameterization to use for numerical solutions. Next, specify a sigma (*σ*), which is a standard deviation of a parameter distribution from which variants can be drawn in multiple runs (not a metric of individual variation within populations). The default choice for *σ* is 0 ^1^. The function `PickTrait` draws a sample from a lognormal distribution for each trait. The next step is to create the initial parameter vector using the parameters picked, and save the names in a separate vector of strings for reference. The Julia code would look something like this –

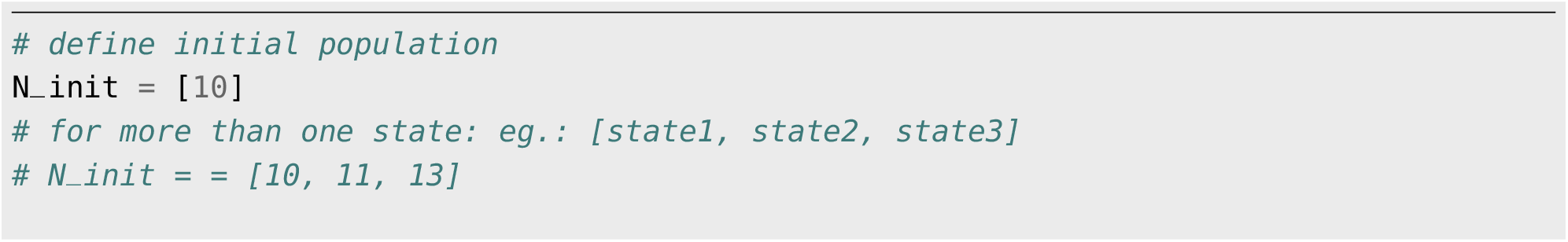

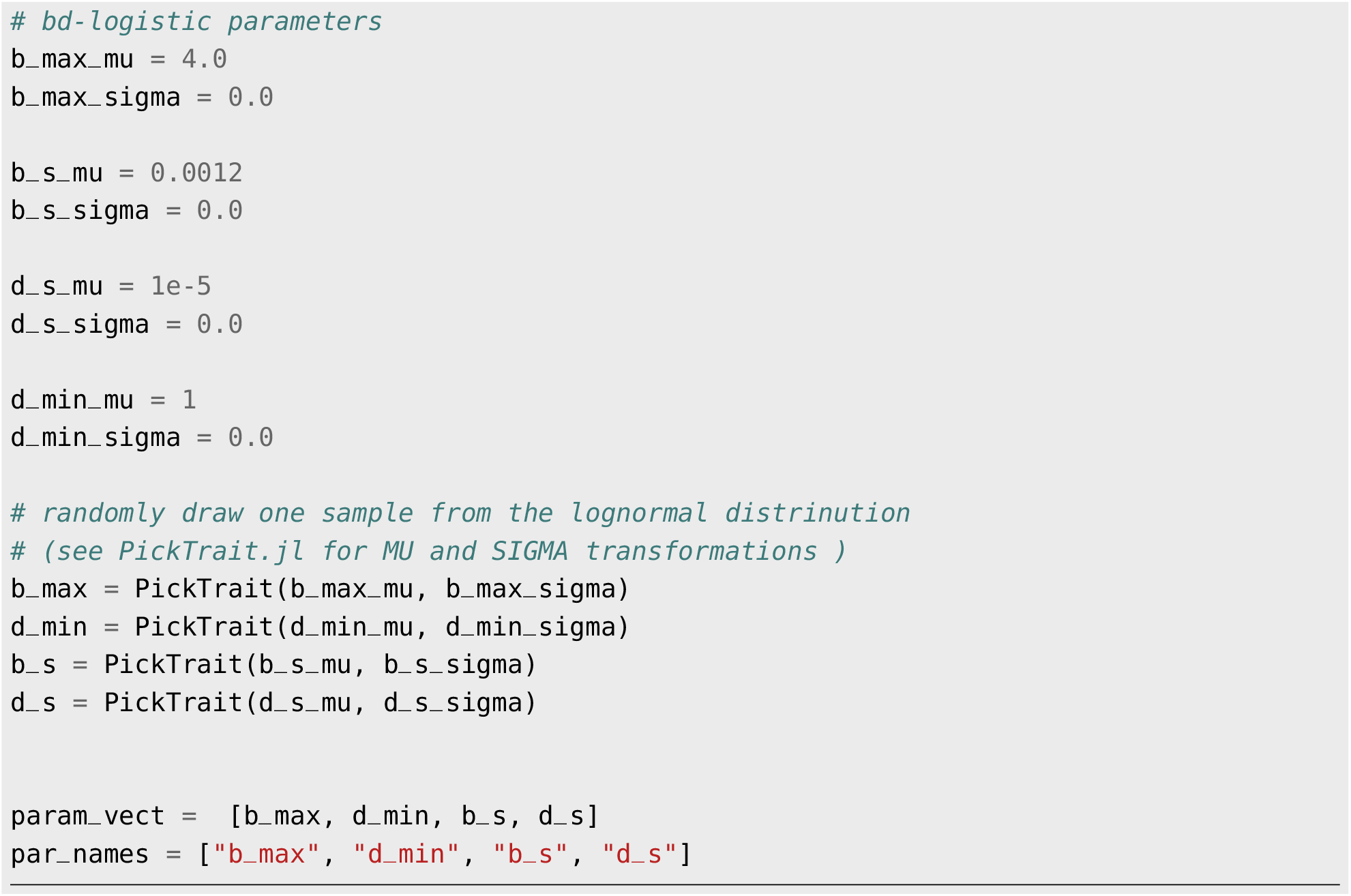

#### 3.3.2 Parameter and Genotype matching

In this block, map the parameters to their states, which means indicate which parameters ‘belong’ to which states (e.g., b_max belongs to state N, not to another state such as a predator). A boolean matrix called `state_par_match` maps the parameters to the corresponding states in the model using a 1 for ‘yes it applies to this state’ or a 0 for ‘no’. The states are represented in rows, and the parameters in columns. Similarly, the `state_genotype_match` matrix indicates which genotypes/clones map onto which state. The algorithm uses these matrices to determine which individuals get which parameters and get genotype-specific information. An example matching-matrix for the b-d logistic model will look something like –

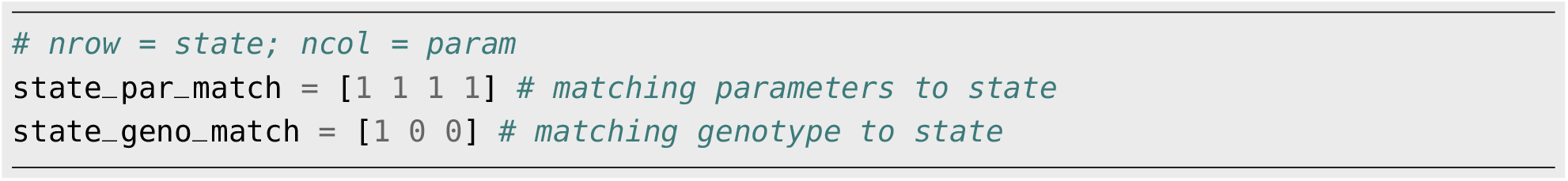

Before wrapping up this section, save the genotype identifier names in another vector. The names can be any kind of useful genotypic handle, such as “Aa”, “Whorled”, or “Genotype 1”. This vector will be used to name the columns in our simulation output.

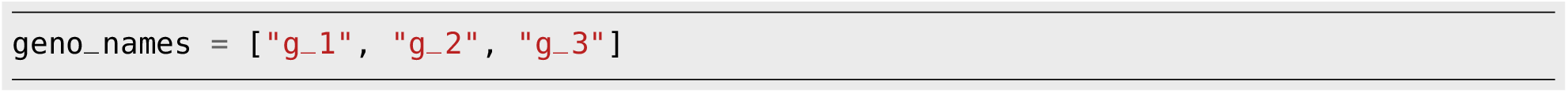

#### 3.3.3 Simulation Design Choices

These are choices an user will make regarding the levels of trait variation, heritability, and conditions of the simulation.

##### GEM versions

In this block, we determine how many different GEM iterations (hereafter referred to as GEM versions) to implement. A common choice is ‘no trait variation’, ‘no evolution’, ‘evolution’, so three different versions that vary by the designation of trait variation and heritability. One may define additional versions depending on other features of interest specific to the model (for example, changing the standard deviation in the initial parameter pick). This part of the code looks like this –

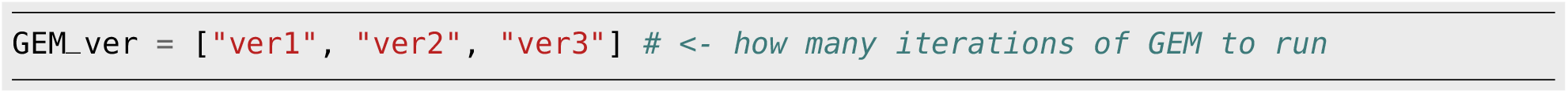

You may choose to have specific descriptive names for the GEM versions like “ver1: no evolution” etc.

##### Heritability and coefficient of variation

GEMs can produce simulations with and without evolution, depending on how the coefficient of variation (CV) and heritability (*h*^2^) are set. For the CV, specify trait variation using a matrix of CV values where the rows correspond to states, the columns to parameters, and stacks to GEM versions. A typical choice includes setting all values in the first stack to zero to indicate no trait variation for GEM version 1, and populating the next stacks with chosen CV levels per parameter per state to introduce variation.

Next, define a matrix for the narrow-sense heritability (*h*^2^) between 0 and 1, corresponding to states in rows and GEM versions in columns. A common choice here would be to have 0s in the first column for traits that are not heritable, but some chosen heritability in the latter columns to activate evolutionary dynamics (i.e., turn evolution ‘on’). An example CV anf *h*^2^ matrix for the b-d logistic model with 3 GEM versions would look something like –

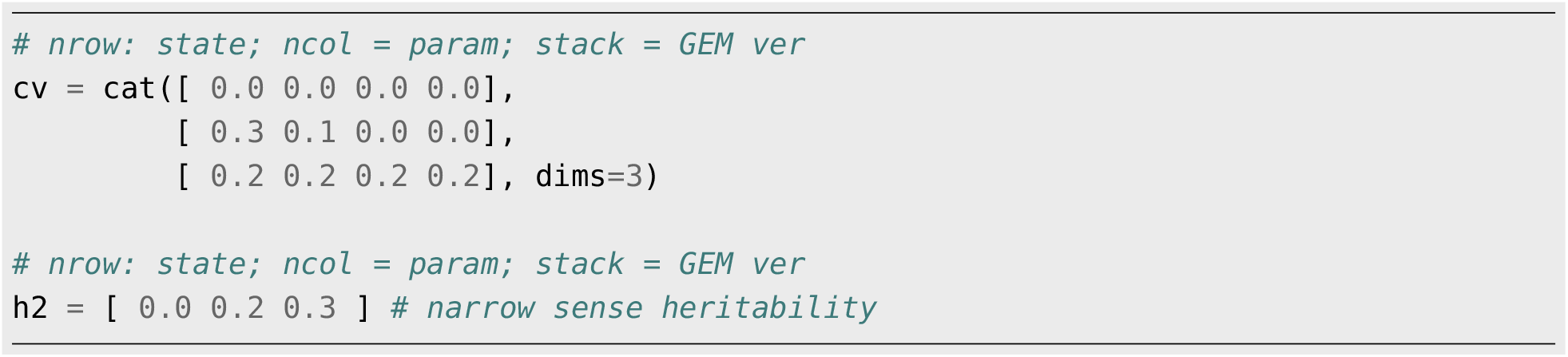

##### Replicates and Time

Next, determine how many independent replicates to run for each version, as well as the total duration of each replicate (*t*_*max*_). Unless all populations go extinct, each replicate simulation will continue until it reaches the maximum user-defined time *t*_*max*_. One challenge in displaying the GEM output efficiently is that each simulation has a different sequence of time steps. As a result, it is tedious to grab percentiles to illustrate the variation in output at given moments in time. To simplify the process and save memory, we log data at standardized time points. The simulations therefore only store abundances and traits as they pass this set of standardized times. The default value is 0.1, that is, a tenth of whatever time unit reflects the parameterization of the model. Memory efficiency can be increased by raising this value and storing fewer points.

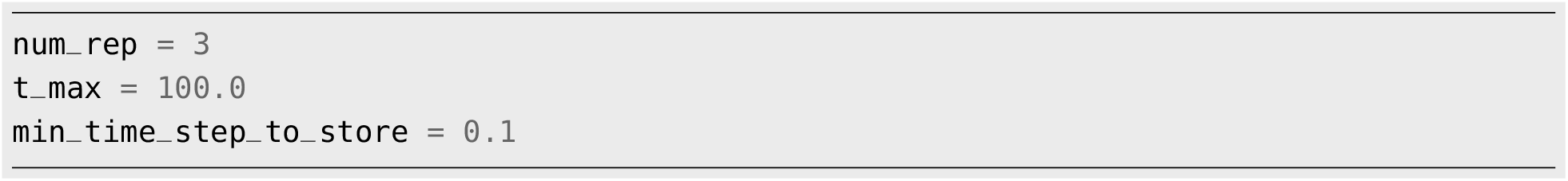

#### 3.3.4 Instantiate model and simulation parameters

The last block in the `model_config.jl` script instantiates the structures created under *model definition* with the parameters and conditions chosen above.

## 4 Run GEM Simulation

With setup complete, the GEM is now ready for execution. This can be done interactively through the REPL or more efficiently using a script (like the provided `GEM_run.jl`). `GEM_run.jl` is used to call scripts to load all required packages, the model definition and configuration files, and all the algorithm functions (Fig 1). Once all of these are loaded into the scope, the GEM simulation can now be executed by calling the `GEM_sim` function from within `GEM_run.jl`. The function accepts arguments for the initial state densities, a vector of model parameters, the simulation design choices, model-to-simulation mapping structures, simulation parameter, and some predefined output container. These arguments are grouped into `struct` (table 1), and should already be instantiated if the `model_config.jl` file is loaded. Setting `verbose = true` will display the time steps between events on the console, which can be useful when screening parameter space. However, a constant stream of output on the console could potentially slow down a large simulation. With `verbose = false`, users will still be able to see the thread ID the simulation function is running on (see section 5.2 for details on parallel computing).

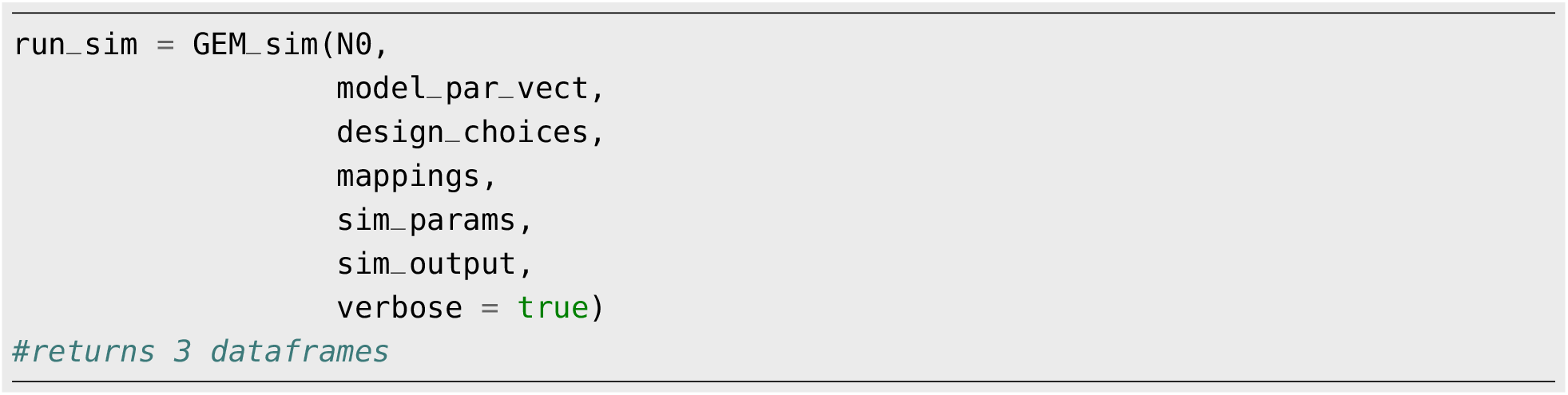

Calling the `GEM_sim` function will execute the main algorithm for all replicates and versions of the model. Upon completion, the function returns three data frames, one for population abundances, one for the trait medians, and one for trait variances. These can be accessed as components of the output with the names `.pop_df` for the population data frame and `.trait_df` for both the trait mean and variance data frames.

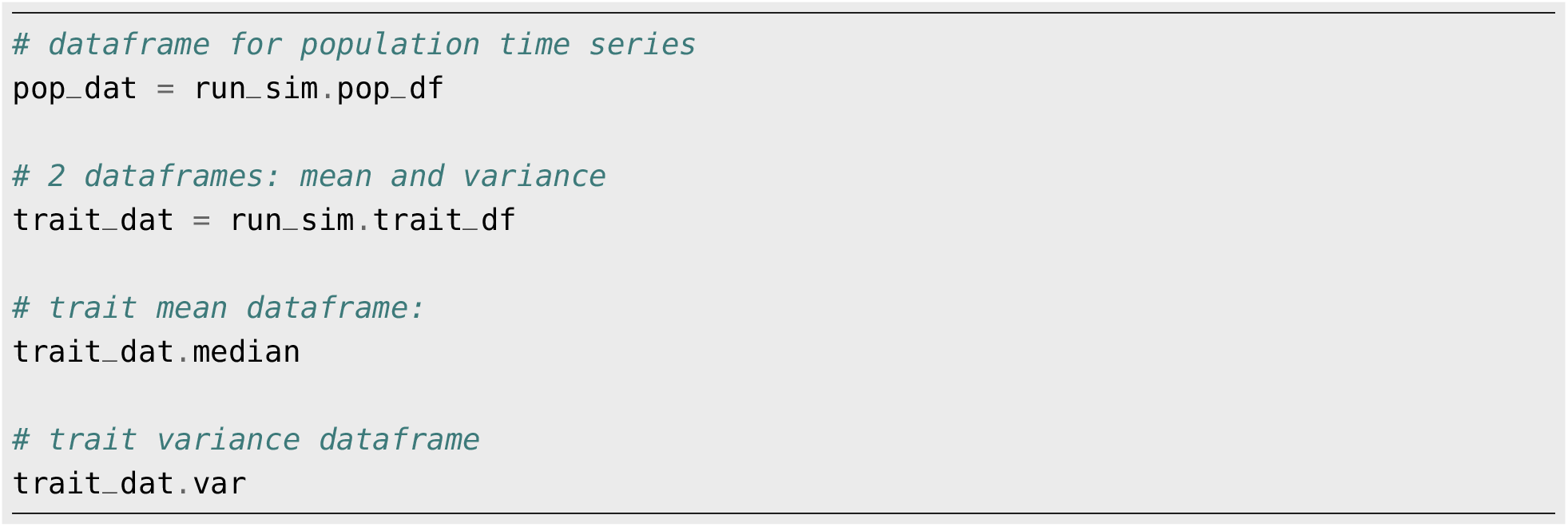

The result data frames can be further analyzed in Julia using `DataFrame.jl` or `Tidier.jl`, or exported as CSV and loaded into your programming language of choice.

The GEM script comes with some introductory plot functions for visualization. All plot functions output every GEM version for each state/trait plotted. Available functions are `Pop_Plot` for population time series plot, `Trait_Plot` for plotting trait time series, and `Geno_Freq_Plot` to plot the genotype frequency. The population plot function (`Pop_Plot`) accepts the population data frame as the first argument, and the state ID as the second argument. The trait time series plot function (`Trait_Plot`) accepts the data frames for trait median and variance, along with the state ID and a string of the trait name of interest, and outputs a timeseries of the trait over time along with the uncertainty. And the genotype frequency plot function accepts the data frame for trait median, along with the state ID, and a string of genotype name to plot. Fig 3 shows the population, trait, and genotype frequency time series for our birth-death logistic model example. We encourage users to add their own plot functions according to the needs of their study and hypothesis.

**Figure 3:**
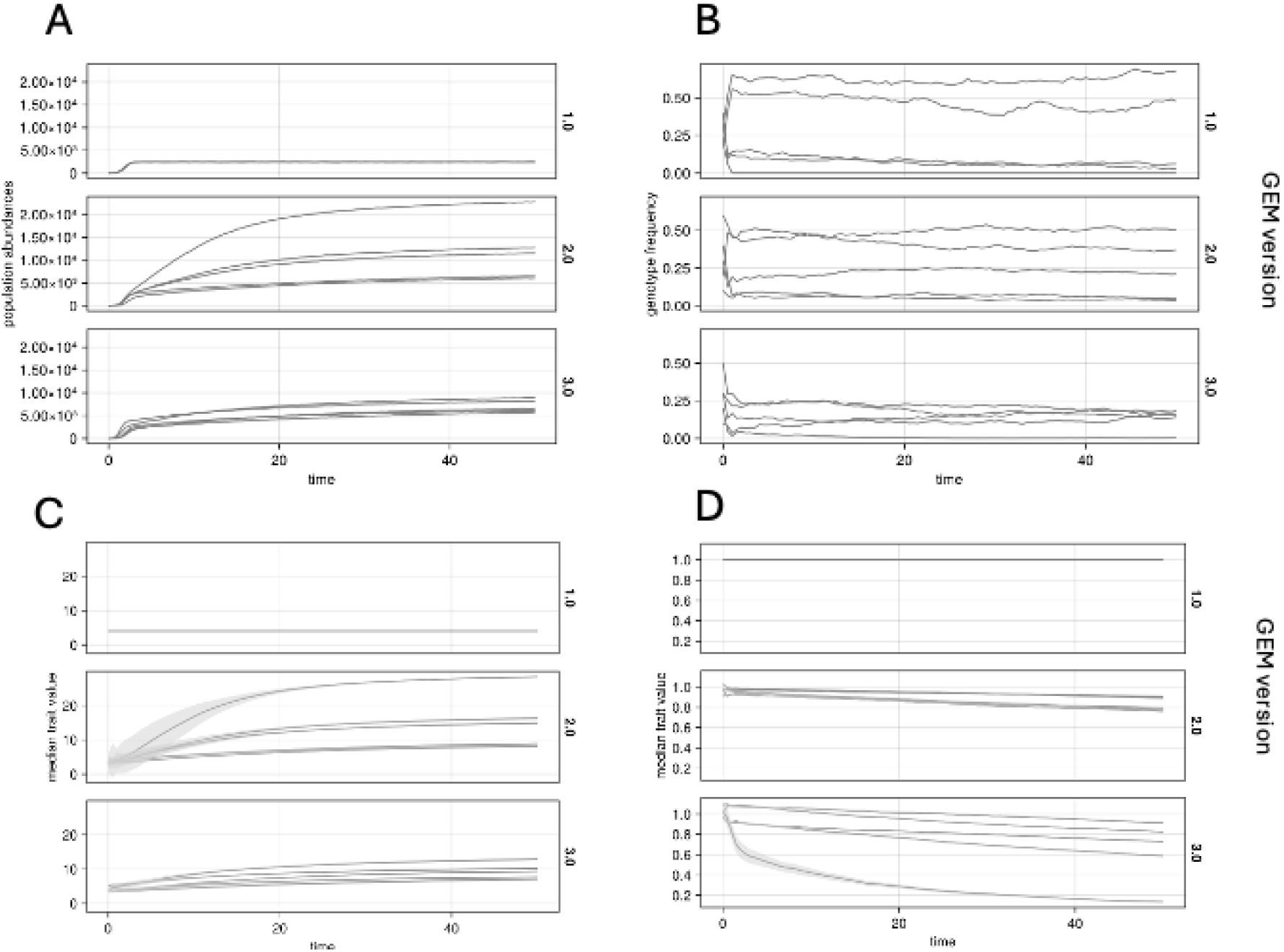
b-d logistic model example output. We ran the model for 50 time units, with three GEM versions, and 5 independent replicates for each version. Initial parameter distribution and starting abundances are the same for each version. GEM versions only differ in their narrow sense heritability *h*_2_, and coefficient of variation (*CV*). Staring conditions are the same for each replicate. `GEM ver1` *h*_2_ = 0 (“no-evolution”); `ver2` *h*_2_ = 0.2, `ver3` *h*_2_ = 0.3. (A): population time series across 3 GEM versions. (B): Genotype frequency for example genotype “g_1”. (C): time series of trait “b_max”. In addition to different *h*_2_ values between versions, b_max also differs in *CV*. `GEM ver2` *CV* = 0.3 and `ver3` *CV*=0.2. (D): time series of trait “d_min”. Here too *CV* value differs between versions. `GEM ver2` *CV* = 0.1 and `ver3` *CV*=0.2.

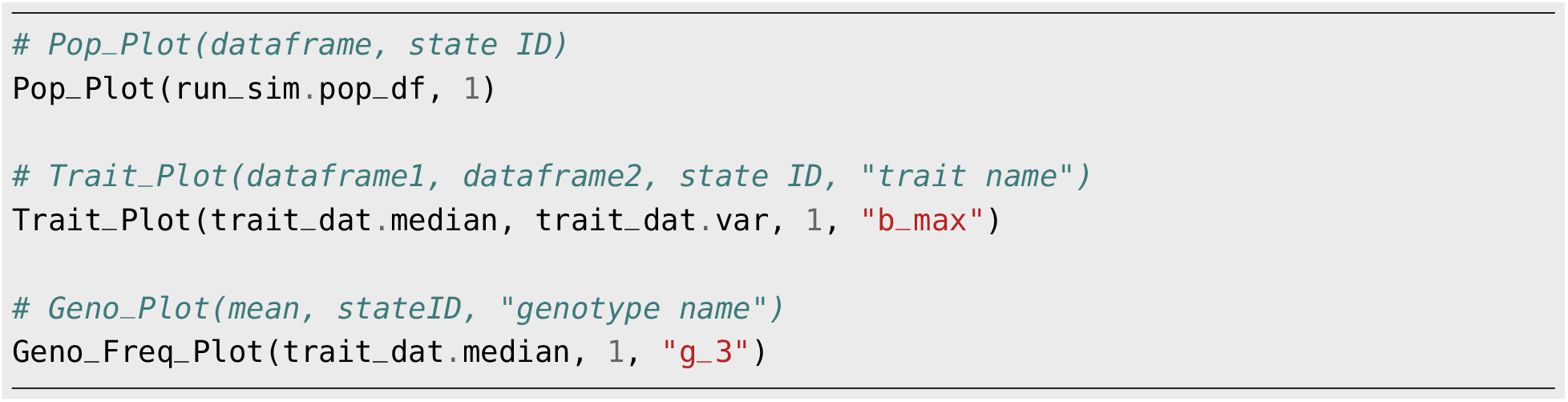

## 5 Additional Considerations

### 5.1 Optional Check on the Deterministic Dynamics

This is an optional step before running the GEM simulation, and it is meant to serve as a check that your model has captured the intended ecological dynamics. Using the defined model and the parameters chosen, one can choose to run a deterministic model using the `DifferentialEquations.jl` (Rackauckas and Nie 2017) package to check the model’s dynamics. This step provides insights into the behavior of the model without any demographic stochasticity and evolutionary processes. Depending on the outcome, initial abundances may be revised or the parameter space adjusted until the desired ecological dynamics are achieved. An example code to assess the deterministic dynamics of the b-d logistic model will look something like –

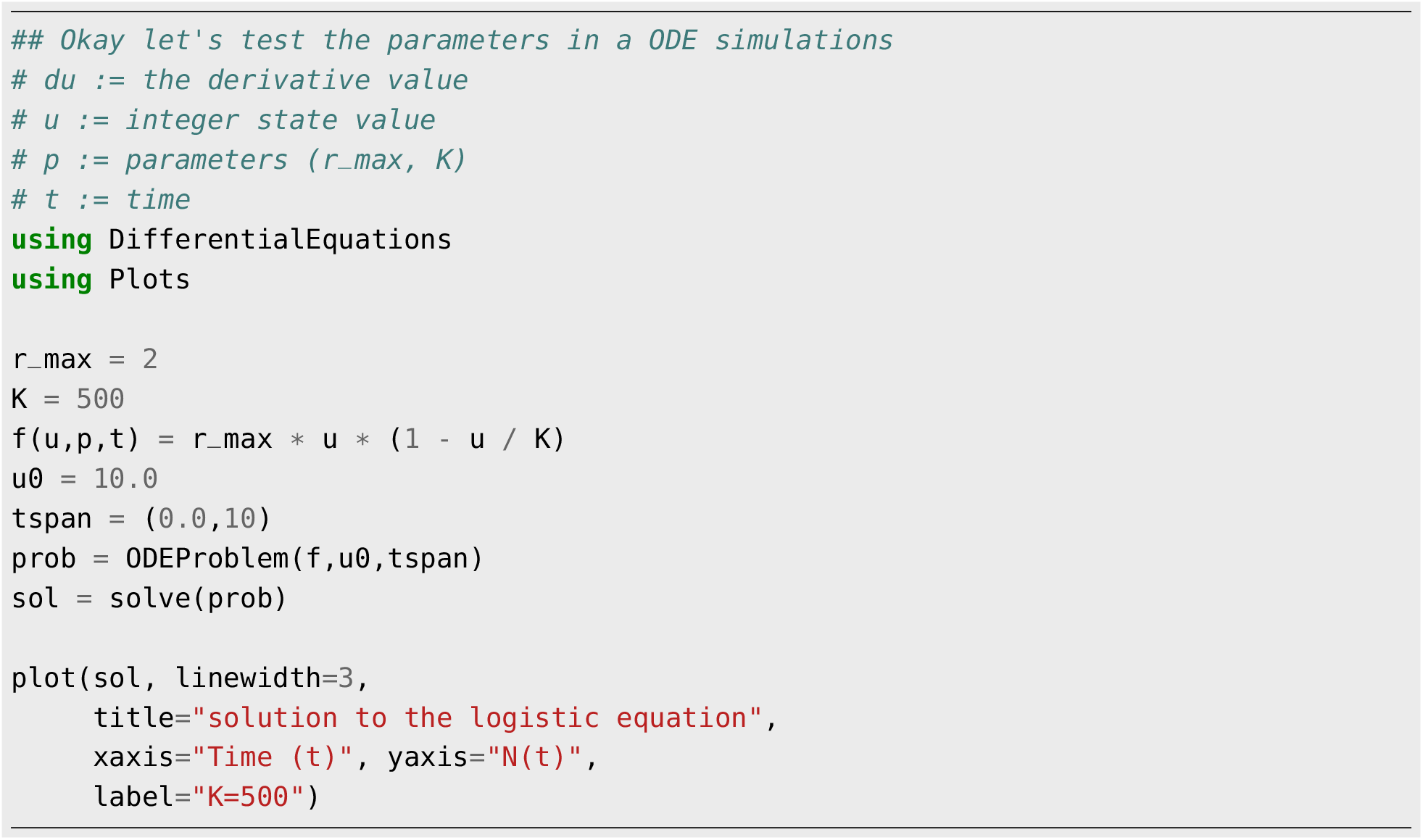

### 5.2 Parallel GEM

GEMs are programmed to utilize multi-threading if the option is made available within the Julia settings. Julia defaults to using one thread. Users can change the default number of threads by accessing the Julia Threads setting in VSCode prior to launching Julia. To set threads to the desired number `n`, go to Settings > search “julia threads” > Julia:Num Threads > Edit in settings.json > `“julia.NumThreads”: n`. The command `Threads.nthreads()` can be used to check the number of available threads, and `Threads.threadsid()` can be used to find the current thread.

### 5.3 Adding New Functions

New functions can be added in two ways. Any new function will need to be loaded into the working environment scope. The function can be loaded into the scope either directly, by adding the line `include(“my_new_function.jl”)` in the `“GEM_run.jl”` file. Alternatively, one can open the `GEM_functions.jl` file and include the new function name here. The latter option keeps the GEM run workflow clean and organized.

### 5.4 GEM as a local package

Advance users may want to turn the core GEM simulation algorithm into a local package that can then be loaded with the `using` command. Users will need to define a module and decide what functions they wish to export. To create a local package, use the `develop` command followed by the path to the directory that contains the package script files. This can be done in the REPL using the package manager (or using the command `Pkg.develop()` outside the package manager)

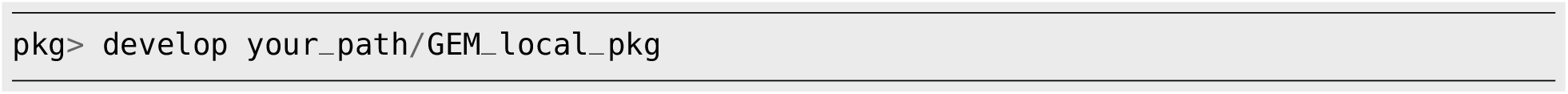

This is a one-time setup step which modifies the `Project.toml` and `Manifest.toml` files that keeps track of all project dependencies. The local package is available for use and can be loaded with the command `using GEM_local_pkg`. This command loads the local GEM package code into the environment ready for use. Changes made directly to the source files will be automatically updated when Julia is restarted.

## 6 Conclusion

The tangled banks of the natural world are home to interacting evolutionary and ecological processes. These banks are often far from equilibrium and are characterized by stochastic birth and death processes among many interacting species. Understanding eco-evolutionary processes in such complex ecological scenarios requires tools that capture the fundamental realities, and GEMs are one such tool. GEMs can be equilibrial but do not need to be; GEMs capture inherent demographic stochasticity as well as the effects of individual trait variation; and GEMs facilitate multi-trait and multi-species evolution. In short, if an ecological model can be written for the system, a GEM can be made for the system. Although originally designed as a quantitative genetics type approach for continuous traits, GEMs now can handle both quantitative traits and discrete traits in the forms of genotypes or strains as well.

This user’s guide is meant to make it easier and faster to build GEMs for new scenarios. We have placed many information containers and design structures into functions that adapt to model configurations chosen by the user, allowing users to focus on model and scenario development to answer new questions. With built-in multi-threading and increased computational speed, we hope the Julia-GEM tool will lead to numerous new insights into eco-evolutionary processes.

## Footnotes

1 When *σ* is >0, the script samples a new parameter value from a Lognormal distribution using the mean and standard deviation provided. This sampled value is then used in one replicate simulation as the mean population parameter. Each subsequent replicate will have another sampled mean. This approach simplifies the process of exploring uncertain parameter space and reduces the need to conduct factorial versions of the model by testing for robustness in outcomes across a wide range of parameters.

